# Machine learning prediction of Antibody-Antigen binding: dataset, method and testing

**DOI:** 10.1101/2021.03.19.435772

**Authors:** Chao Ye, Wenxing Hu, Bruno Gaeta

**Affiliations:** School of Computer Science and Engineering, the University of New South Wales, Sydney NSW 2052

**Author notes:** Corresponding Author Chao Ye.

## Abstract

DNA sequencing technologies are providing new insights into the immune response by allowing the large scale sequencing of rearranged immunoglobulin gene present in an individual, however the applications of this approach are limited by the lack of methods for determining the antigen(s) that an immunoglobulin encoded by a given sequence binds to. Computational methods for predicting antibody-antigen interactions that leverage structure prediction and docking have been proposed, however these methods require knowledge of the 3D structures.

As a step towards the development of a machine learning method suitable for predicting antibody-antigen binding affinities from sequence data, a weighted nearest neighbor machine learning approach was applied to the problem. A prediction program was coded in Python and evaluated using cross-validation on a dataset of 600 antibodies interacting with 50 antigens. The classification predicting accuracy was around 76% for this dataset. These results provide a useful frame of reference as well as protocols and considerations for machine learning and dataset creation in this area.

Both the dataset (in csv format) and the machine learning program (coded in python) are freely available for download.

## 1. Introduction

DNA sequencing technologies are providing new insights into the immune response by allowing the large scale sequencing of rearranged immunoglobulin gene present in an individual [1-2]. However the applications of this approach are limited by the lack of methods for determining the antigen(s) that an immunoglobulin encoded by a given sequence binds to. Individual immunoglobulins can be tested experimentally at significant cost, however the large scale characterization of binding properties from sequence data is currently impossible.

Computational methods for predicting antibody-antigen interactions that leverage structure prediction and docking have been proposed [3]. However these methods require knowledge of the 3D structures. Direct prediction of antibody-antigen interactions from protein sequences remains an open problem.

Machine learning has had some success in predicting antibody interactions in other cases, such as mCSM-AB[4] and ADAPT[5]. mCSM-AB is a web server for predicting changes in antibody-antigen affinity upon mutation, using graph-based signatures. Assisted Design of Antibody and Protein Therapeutics (ADAPT) is an affinity maturation platform interleaving predictions and testing that has been previously validated on monoclonal antibodies (mAbs).

As a step towards the development of a machine learning method suitable for predicting antibody-antigen binding affinities from sequence data, weighted nearest neighbor and Random forest machine learning approaches have been applied to the problem. A prediction program was coded in Python and evaluated using cross-validation on a dataset of 600 antibodies interacting with 50 antigens. The classification predicting accuracy was around 76% for this dataset.

These results provide a useful frame of reference as well as protocols and considerations for machine learning and dataset creation in this area. The method is still limited due to the scarcity of training data but its usefulness for large scale prediction should increase as more antibody-antigen data become available. The ability to predict an antibody’s binding will allow leveraging large scale immune receptor sequencing to allow understanding of how antibody-antigen binding specificities vary across the board within an organism over time, under a range of conditions, and between individuals and populations.

## 2. Method

### 2.1 Dataset

Due to the scarcity of suitable antibody-antigen pairs, computational docking was used to generate some of the training and testing dataset. The Cluspro and Rosetta online web servers [6-9] were used to create a dataset of paired antibody-antigen complexes for machine learning. Both ClusPro and Rosetta were used for protein-protein molecular docking [6-9]. Rosetta uses the SnugDock algorithm [10-12]. Swiss PDB viewer [13] was used to examine the resulting protein complex structures.

Fifty antibody-antigen complexes were selected randomly from the Protein DataBank (www.rcsb.org). The antibody-antigen complexes were separated using Perl script to produce pdb files as well as sequences for antibodies and antigens. CDRs were located using the Rosetta antibody modeling web server. Antigens were docked with a range of antibodies using Cluspro. Ten to Fourteen antibodies were docked with each antigen in order to find the best orientation. Those resulting complexes were submitted to the Rosetta web server in order to calculate the best interface score. Altogether, 50 antigens were docked with 600 antibodies. An example of resulting complex is shown in figure 1.

**Figure 1:**
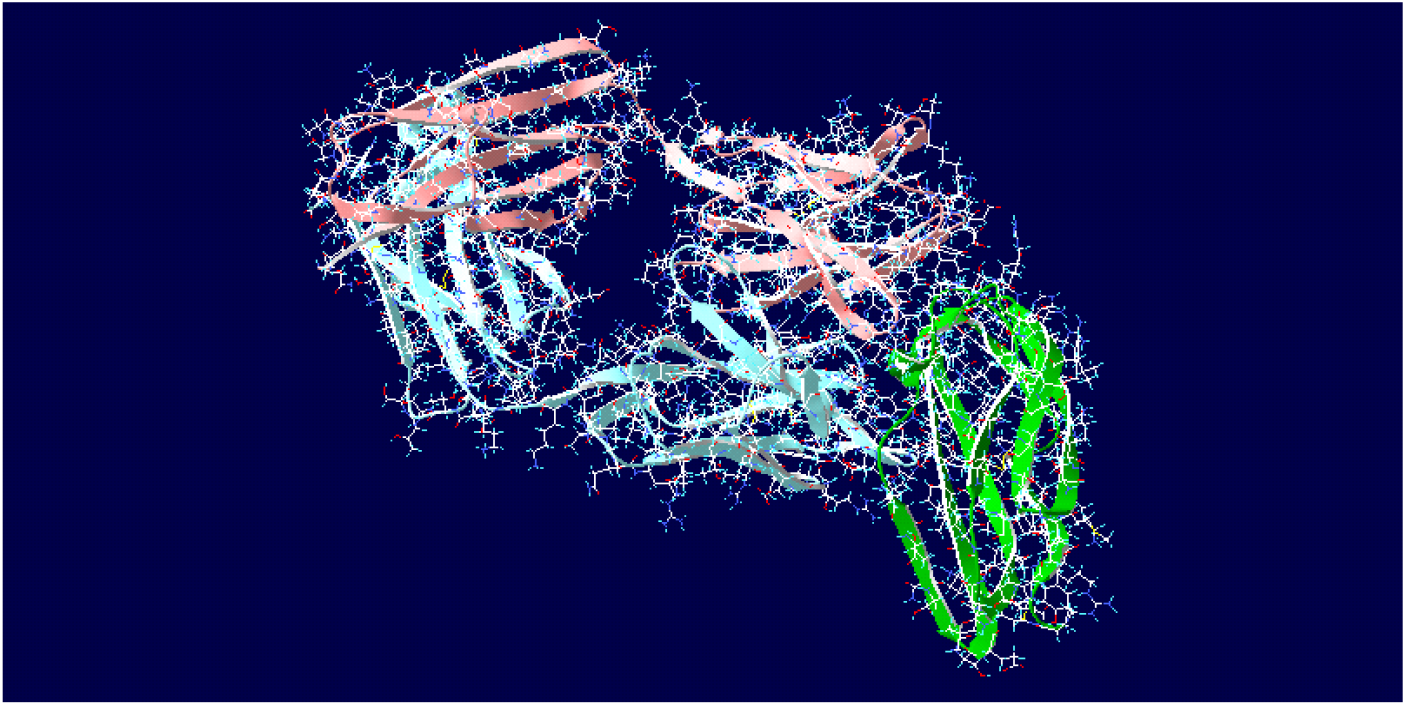
Docking example of 3s35 complex. (Docking results YES Docking Best interface Score -0.876)

The Rosetta interface scores were used as estimates of the binding affinity, as input for machine learning. Good binding affinity was set to correspond to interface scores lower than -8.5. Otherwise, complexes were considered to show less than good binding affinity. In the case of scores between -8.0 and -9.0, the docking clusters and positions were examined visually using SwissDock. If the top 10 models had their antibody and antigen in similar relative positions and the structures showed sensible interaction patterns, the pairs were assumed to have good binding affinity.

Rosetta interface scores have been used previously as classifier to determine binding affinity based on docking results, for example in an antibody-antigen cross reactivity study [14].

In order to further analyze the docking results, the IEDB analysis tool http://tools.iedb.org/bcell/ was used for B cell epitope prediction. The predicted B cell epitopes were recorded as part of the machine learning dataset.

Additional data were extracted from the CoV-AbDab [15] database of antibodies against coronaviruses, including SARS-CoV2, SARS-CoV1, and MERS-CoV. Data (2674 rows) were obtained from CoV-AbDab on 14/02/21. After filtering of incomplete data, 2031 rows remained, with each row corresponding to an antibody. Since a row may contain information about an antibody’s interactions with multiple antigens, the data was further split into multiple rows, each containing information about the interaction between an antibody and an antigen and marking whether it can bind. This produced a total of 4441 rows of data, including 2850 positive samples and 1591 negative samples.

Among the dataset, 1057 antibodies were identified to bind to the receptor binding domain (RBD) of the spike protein for SARS-CoV2, SARS-CoV1 and MERS-CoV. This generated 1675 rows of data, which included 1265 positive samples and 410 negative samples.

Additional features for machine learning were generated using the Bachem peptide analysis tool https://www.bachem.com/service-support/peptide-calculator/ to calculate the Isoelectric point and net charge at neutral pH (7.0) for each of the CDR features. These CDR features were included in the machine learning dataset for random forest machine learning.

The resulting dataset can be downloaded as supplementary data and is structured under the following column headings:

- H chain CDR1 sequence
- H chain CDR2 sequence
- H chain CDR3 sequence
- L chain CDR1 sequence
- L chain CDR2 sequence
- L chain CDR3 sequence
- Hydro of L CDR1
- Pl of L CDR1
- Hydro of L CDR2
- Pl of L CDR2
- Hydro of L CDR3
- Pl of L CDR3
- Hydro of H CDR1
- Pl of H CDR1
- Hydro of H CDR2
- Pl of H CDR2
- Hydro of H CDR3
- Pl of H CDR3
- Antigen Epitope
- Rosetta Docking score
- Antigen
- Docking result

### 2.2 Machine Learning Method Overview

A weighted K-nearest neighbors (k-NN) classification algorithm for predicting antibody-antigen binding affinity was implemented in Python. The program can be downloaded as supplementary data.

Weighted K-nearest neighbors (k-NN) method:

Neighbors were determined using string distances between the CDR1, CDR2 and CDR3 amino acid sequences of different antibodies. Weights were calculated from distances so that nearer neighbors were considered to have more weight as detailed below.

For every antigen, the binding affinity was predicted using the k-NN method for between 10 and 15 randomly selected antibodies. K nearest neighbors were identified by comparing their string distances in order to classify antibody-antigen pairs as “good affinity” or “low affinity”. The predicted results were then compared with the actual classification obtained from docking. In order to ensure the k nearest neighbors pairs only included pairs with same antigen, a fixed penalty of 1000 was added to the distances between antibody-antigen pairs involving different antigens.

As antigens and epitopes vary significantly, antibody CDR distances were selected as the initial starting point for machine learning. The program started learning from 10-15 antibodies interacting with one antigen, then looped to include up to 40 antigens.

This approach makes antibody-antigen binding affinity a computational problem suitable for machine learning.

### 2.3 Antibody Distance calculation for the K-nearest neighbors method

#### String distance calculation

The similarity between antibodies was measured by comparison of their CDRs. Each antibody has a heavy chain and a light chain and each contains 3 CDRs. The distance between two antibodies was calculated as the Euclidian distance between their CDR distance vectors as shown in Eq. 1. The Python code is given in Box 1.

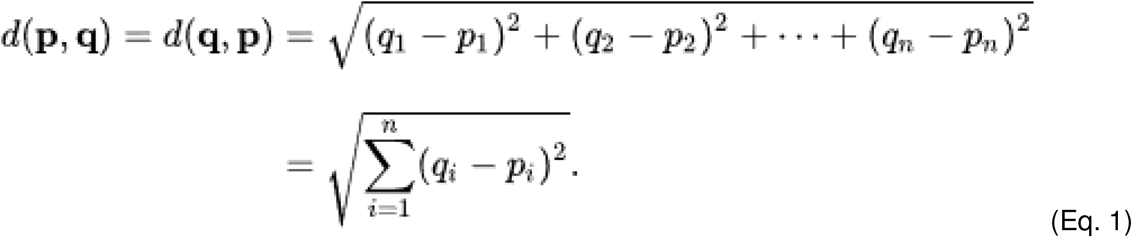

(qi-pi) = string distance between Cdri of antibody q and Cdri of antibody p

##### Box 1

**Python code for Euclidian distance calculation**

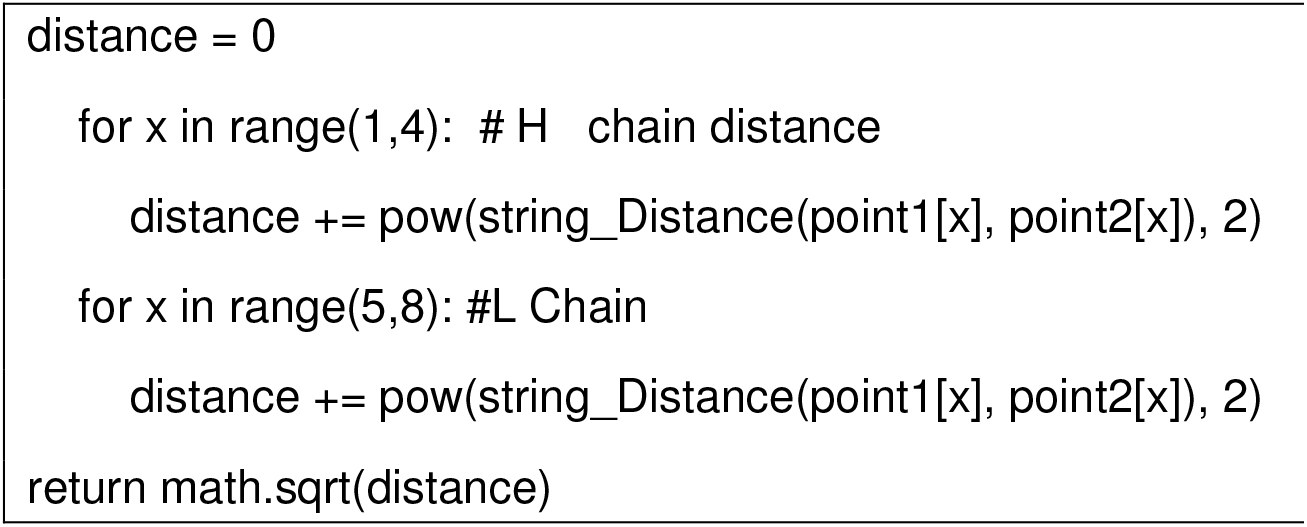

Two different CDR distance calculation methods were tested and compared:

#### 1. Levenshtein distance based on amino acid identity

Pairs of equivalent CDRs were compared with each other using their Levenshtein string distance [16] as shown in Eq. 2

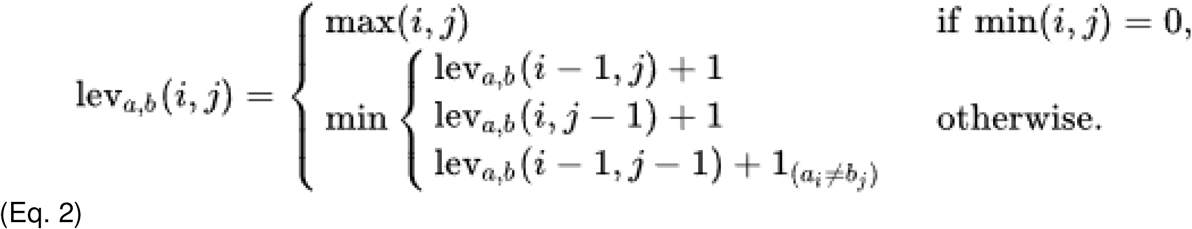

**Cost =0 for ai=bi**
**Cost =1 while ai=/=bi**

#### 2. Levenshtein distance with Blosum62

The Levenshtein distance only considers amino acid identity when comparing sequences. A more biologically significant distance measure needs to take into account the different properties of amino acids, which means that some amino acid substitutions are more likely to be accepted in an interaction than others. The BLOSUM62 substitution matrix [17] was used as a proxy for amino acid similarity in the Levenshtein distance calculation. Although the BLOSUM matrices were designed to reflect evolutionary conservation, they can provide an estimate of similarity in interaction potential [18].

Levenshtein distance was calculated as per Eq. 2 but using the following cost function:

**For ai=bi, Cost =0**

**For ai=/= bi, Cost = Dij =**| **Sij - (Sii + Sjj)/2**| **/2**

where Sii, Sjj, and Sij are obtained from the BLOSUM62 matrix.

### 2.4 Random forest Machine learning algorithm

A random forest machine learning algorithm incorporating the previous k-NN results was also used for predicting antibody-antigen binding classification.

Isoelectric point and Net charge at neutral pH (7.0) for each CDR were used as additional features in addition to the BLOSUM62-derived CDR distance for training the random forest. Binding was predicted based on combining the votes from each of the features, with each of the individual features contributing one vote according to the nearest neighbor prediction based on that feature.

## 3 Results, testing and discussion

### Classification results

The antigen-antibody binding classification methods were evaluated using leave-one-out cross-validation and are summarized in Table 1. For the k=2 nearest neighbors method with Levenshtein distance based on sequence identity, the accuracy was 69%. A slight improvement (accuracy of 72%) was observed when using the BLOSUM 62 matrix to calculate the Levenshtein string distance.

**Table 1:** Comparison of antigen-antibody binding prediction approaches: classification accuracy (binding/non-binding) for the antibody-antigen pairs in the dataset was estimated using leave-one-out cross-validation.

| Method | 2 nearest neighbor with string distance | 2 nearest neighbor with string distance and Levenshtein distance | Random forest (best) |
| --- | --- | --- | --- |
| Accuracy | 69% | 72% | 76% |

Different k values were also evaluated with the Levenshtein distance based on BLOSUM 62. A value of k=2 provided the best accuracy. For k=1 nearest neighbor, accuracy was 71%. For k=3, classification accuracy dropped to 68.9%.

For the random forest predictions, votes were taken as classification prediction results. The accuracy was highest when the whole forest was considered, in which case each feature contributed to the classification results. A slight improvement (accuracy 76%) was observed when all 13 features (the Levenshtein string distance, and the isoelectric point and net charge at neutral pH (7.0) for each CDRs) took part in the final votes.

### Regression to predict docking score

The nearest neighbor method (k=2) was also evaluated by examining its ability to predict the docking score by regression using leave-one-out validation. Actual and predicted scores are compared in figure2. The BLOSUM 62-based method resulted in a slight improvement over the sequence identity-based method. Reducing the number of rows used to calculate the mean standard error to 200 did not affect the values much (MSE = 2.63).

**Figure 2:**
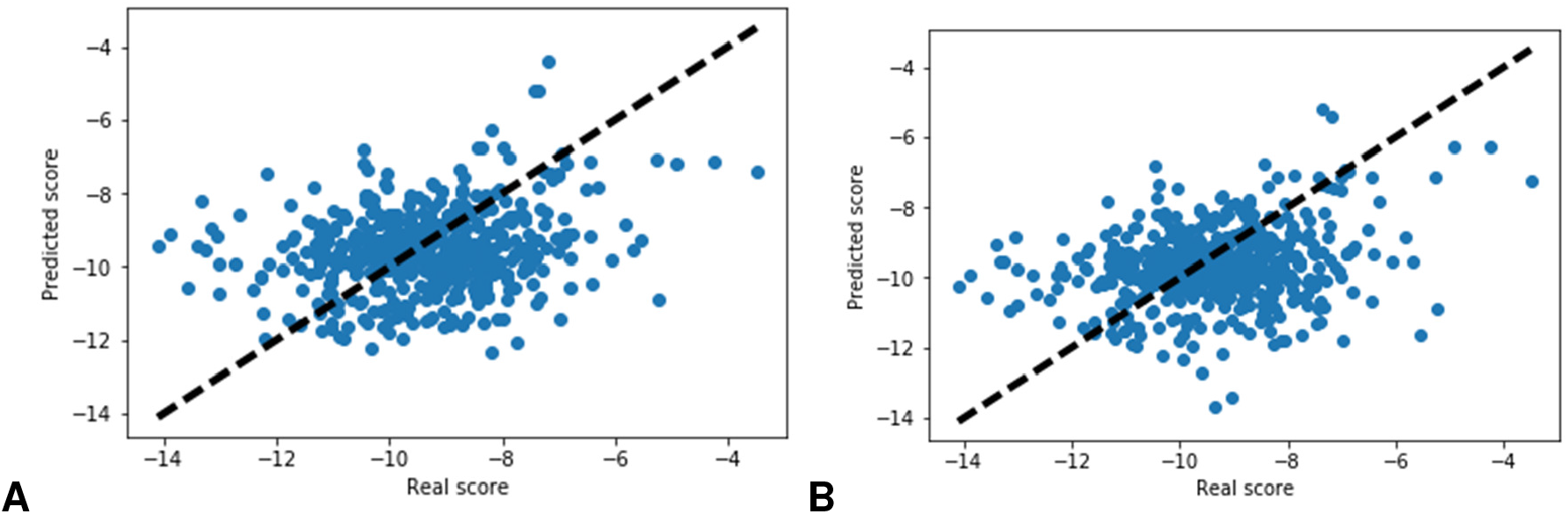
Comparison of docking scores predicted by nearest neighbor based on Levenshtein Distance (k=2) with actual scores. A. Prediction based on sequence identity. B. Prediction based on BLOSUM 62 matrix. The BLOSUM 62 prediction resulted in a slight improvement (Mean Standard Error = 2.71, 500 rows) over the sequence identity-based prediction (MSE = 2.72, 500 rows)

Unlike the classification, increasing the value of k resulted in a slight improvement in the prediction of docking scores. With k=4, the mean standard error was 2.13 based on 200 rows.

## Discussion

A 500 rows dataset of antibody-antigen pairs was created for training and evaluation of a new program for predicting antibody-antigen interaction parameters using machine learning. Subsets of this dataset (200, 300, and 400 rows) were tested during the data collection process.

Classification accuracy was quite consistent around 77% across all these subsets. While the dataset is limited, it provides a good starting point for the development of the method, and validates the use of this approach for prediction of antibody-antigen binding affinity as more data become available.

The KNN nearest neighbor method was chosen as machine learning method. The best prediction results were obtained with 2 nearest neighbors (K=2). Random forests were also used that incorporated sequence distance as well as chemical properties of the CDRs (pI and hydrophobicity). The best prediction results (accuracy of 76%) were obtained with a full forest which used all the provided features.

In the absence of large amounts of experimental data on antibody-antigen binding affinities, Rosetta interface scores along with top 10 binding positions were used to determine the classification for binding affinity. While this is unlikely to provide a full representation of the problem, it provides a dataset suitable for comparing a range of approaches. The method will certainly improve as larger datasets become available.

Around 20% of the method’s predictions were inaccurate. These errors mostly happened with some large antigens. The docking results for these antigens were further examined. The decreased accuracy is likely to be the result of conformational flexibility in the larger antigens, the presence of multiple epitopes, and the increased number of discontinuous epitopes relative to smaller antigens

## 4 Conclusions

We have created a training and test data set of 600 antibody-antigen complexes using a combination of structure modelling and computational docking using Rosetta.

We also developed weighted nearest neighbor and random forest approaches to predict antibody-antigen binding based on sequence data. These machine learning procedure can use classification to predict cognate antigen. When using k-NN based on sequence distance, regression can also be used to predict Rosetta binding scores.

Leave-one out cross-validation testing yielded an accuracy of 72 % for classification results based on the 2 nearest neighbors. The prediction accuracy was around 67% -76% when varying the number of nearest neighbours. The best prediction results (accuracy of 76%) were obtained with a full forest which used all the provided features.

